# Age-related macular degeneration-like phenotypic features develop at the early ages of Cxcr5/Nrf2 double knockout mice: An accelerated AMD model

**DOI:** 10.1101/868851

**Authors:** Hu Huang, Anton Lennikov

## Abstract

Age-related macular degeneration (AMD) is a leading cause of blindness for older adults. The aim of this study is to develop an accelerated mouse model of AMD and characterize its phenotypic features. Cxcr5 knockout (KO) mice and Nrf2 KO mice were bred to create Cxcr5/Nrf2 double knockout (DKO) mice. AMD-like features in Cxcr5/Nrf2 DKO mice were compared with those in CXCR5 KO mice and C57BL6 wild-type (WT) controls. The assessment included fundus and optical coherence tomography (OCT) imaging, periodic acid-Schiff (PAS) and immunofluorescence staining of retinal pigment epithelium (RPE)–choroid flat mounts and sections. Stained samples were imaged with fluorescent microscopy, and Western blots were used to monitor protein expression changes. The staining of cleaved caspase-3, peanut agglutinin (PNA) lectin, and MAP2 was performed to assess the presence of retinal degeneration and cell apoptosis. Quantification with statistical analysis was performed with Graphpad software. The 2- 4-, and 6-month-old DKO mice exhibited increased hypopigmented spots on fundus and sub-RPE abnormalities on OCT as compared to the Cxcr5 KO mice, and C57BL6 WT controls. Aberrant RPE/sub-RPE depositions and increased Bruch’s membrane (BM) thickness were demonstrated by PAS-stained sections. The DKO mice had strong autofluorescence (A2E) and increased RPE/sub-RPE depositions of IgG and AMD-associated proteins (β amyloid, Apolipoprotein E, complement 5b-9, and αB-crystallin). The protein expression of AMD-associated proteins and Transmembrane Protein 119 (TMEM119) microglia marker were upregulated at the RPE/BM/choroid complex of DKO mice. The adult DKO mice underwent accelerated retinal degeneration and cell apoptosis compared to the KO and the WT mice. Together, the data suggest that the Cxcr5/Nrf2 DKO mice develop significant AMD-like characteristics at an early age and may serve as an accelerated AMD model.

**Sumary Statement:** A new animal model is developed to mimic early AMD characteristics in adult mice

## Introduction

Age-related macular degeneration (AMD) is a complex disease, as exemplified by its association with various genetic polymorphisms and environmental risk factors and its heterogeneous clinical manifestations and pathological features, including the early hallmarks of aberrant sub-retinal pigment epithelium (RPE) and sub-RPE, and sub-retinal deposits such as drusen.(Sarks 1980, Hageman, Luthert et al. 2001, Anderson, Mullins et al. 2002, Curcio 2018) RPE death and photoreceptor degeneration are involved in geographic atrophy (GA), or the “dry” form of AMD,(Datta, Cano et al. 2017) whereas sub-retinal invasion of choroidal vessels or choroidal neovascularization (CNV) is a feature of the “wet” form of AMD.(Bhutto and Lutty 2012) These pathological characteristics are the consequence of both genetic variant predispositions and environmental risk factors. Among the cloned and mapped genes that may predispose individuals to AMD are complement factor H (CFH)(Edwards, Ritter et al. 2005, Haines, Hauser et al. 2005, Klein, Zeiss et al. 2005), apolipoprotein E (APOE) (Klaver, Kliffen et al. 1998), C-X3-C motif chemokine receptor 1 (CX3CR1),(Tuo, Smith et al. 2004, Schaumberg, Rose et al. 2014) age-related maculopathy susceptibility 2 (ARMS2), and HtrA serine peptidase 1 (HTRA1).(Edwards, Chen et al. 2008, Cho, Wang et al. 2009) The known environmental risk factors for AMD include cigarette smoke, blue light exposure, advanced age, high-fat diet, and others. Interactions between multiple AMD risk factors may heighten the pathological processes that damage the photoreceptors and RPE, and thereby resulting in the initiation and progression of AMD.

Reliable and reproducible animal models of AMD are essential for deciphering disease etiopathogenesis and developing effective therapies. Despite the absence of a macula in the mouse retina, mice are widely used to create AMD models, primarily because mouse strains can be easily genetically manipulated and are cost-effective for experimental studies. A number of mouse strains developed in recent years recapitulate some of the essential characteristics of human AMD, such as RPE pathologies, sub-RPE deposition, and RPE/photoreceptor death, including mice deficient in antioxidant factor genes, such as superoxide dismutase (SOD1),(Imamura, Noda et al. 2006) in nuclear factor erythroid 2-related factor 2 (NRF2).(Zhao, Chen et al. 2011) in chemokine receptor, such as C-C motif chemokine receptor 2 (Ccr2)(Ambati, Anand et al. 2003) and Cx3cr1(Combadiere, Feumi et al. 2007) genes. Other experimental strains developed for AMD studies include mice immunized with carboxyethylpyrrole protein (CEP)-adducted protein or antibodies,(Hollyfield, Bonilha et al. 2008, Hollyfield, Perez et al. 2010) mice with apolipoprotein E (ApoE)(Malek, Johnson et al. 2005) mutations, and mice that lack RPE-derived soluble vascular endothelial growth factor (VEGF).(Saint-Geniez, Kurihara et al. 2009)

Recently we characterized the eye fundus phenotypes of aged C-X-C motif chemokine receptor 5 (Cxcr5) knockout mice and found features emblematic of AMD, such as sub-RPE deposits with drusen appearance and RPE degeneration (which are associated with increased inflammation and immune dysregulation), increased inflammatory marker cyclo-oxygenase-2, and microglia activation markers ionized calcium-binding adaptor molecule 1 (Iba1) and arginase 1 (Arg-1), as well as AMD-associated proteins such as β amyloid, complement 3d (c3d), and αB-crystallin (all are deposited at RPE/sub-RPE space and their protein levels were escalated in the RPE/BM/choroid complex protein extracts of the aged Cxcr5^-/-^ mice compared to the same age wild type controls).(Huang, Liu et al. 2017) Moreover, the increased Iba1 and β-amyloid are localized in the sub-retina and in sub-RPE, indicative of a potential role in promoting abnormal sub-RPE depositions and AMD pathogenesis.(Lennikov, Saddala et al. 2019)

A significant challenge to delineating the etiology and pathophysiology of AMD in animal models is that AMD-like features generally develop in advanced-age animals, necessitating a significant amount of lead time for AMD to develop and thus increasing the costs of research. To circumvent this issue, we sought to develop an accelerated animal model of AMD, in which AMD-like features develop at an early age, by combining the two well-characterized pathological factors of oxidative stress and inflammation into one mouse strain. We chose the Cxcr5 knockout (KO) mice and Nrf2 KO mice because they develop increased inflammation and oxidative stress, thus resulting in the AMD-like features in their respective aged animals.

## Results

### Abnormal Fundus and Sub-RPE Deposits in the adult Cxcr5/Nrf2 double knockout mice

Fundus findings in the 4-month-old Cxcr5/Nrf2 DKO female mice were compared with those in the age- and gender-matched wild-type (WT) and Cxcr5 KO controls. As expected, the WT animals demonstrated features of a healthy fundus (Fig. 1A) and the CXCR5 KO mice only had a few hypopigmented spots (Fig.1B). In contrast, numerous hypopigmented spots were visualized in the fundus of Cxcr5/Nrf2 DKO mice (Fig.1C). The fundus observations were further confirmed by OCT imaging, which demonstrated sub-RPE abnormalities that corresponded to hypopigmented areas observed by fundus imaging (Fig. 1D-F) and the 10 distinct retinal layers from the ganglion cell layer (GCL) to the choroid of the mid-peripheral retina in both KO and WT controls (Fig. 1G-I). Quantification indicated that the numbers of hypopigmented spots on the fundus were significantly higher in the DKO mice than in the KO mice and in the WT controls (Fig. 1J). The sub-RPE abnormalities on the OCT graphs were also significantly higher in the DKO mice than in the KO mice and in the WT controls (Fig. 1K).

**Figure 1.**
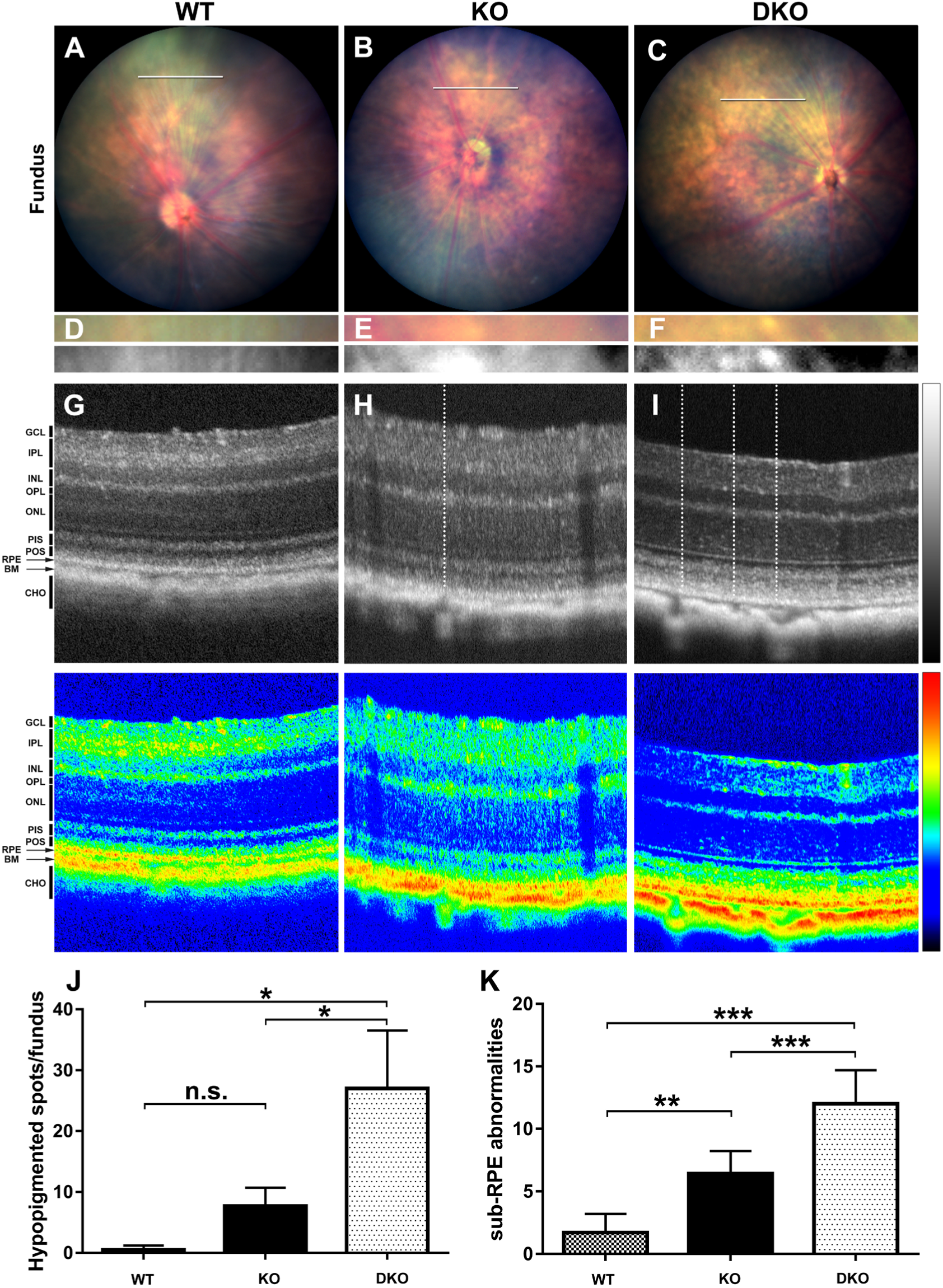
Fundus and OCT images in 4-month-old C57BL/6 WT, CXCR5 KO, and CXCR5/NRF2 DKO mice. The representative fundus images C57BL/6 WT (A), CXCR5 KO (B); CXCR5/NRF2 DKO (C), white bars in the fundus images indicate the area of the corresponding OCT scan. The enlarged portion of the fundus image corresponding to the OCT scan area and contrast-enhanced monochrome version of the same image demonstrate hypopigmented spots (D-F). The representative OCT images (top) and the corresponding OCT heatmaps (bottom) (G-I). White dotted lines map the fundus features in the enlarged high-contrast portions of the fundus images to the abnormal sub-RPE deposits. The quantification of hypopigmented spots in the fundus images (J). The quantification of sub-RPE abnormalities in the OCT data (K). The spots numbers on fundus images and the sub-RPE abnormalities were counted by “masked” observer and averaged from images acquired from 4 animals per group (n = 4). Retinal layers were denoted as follows. GCL: ganglion cell layer. IPL: inner plexiform layer. INL: inner nuclear layer. OPL: outer plexiform layer. ONL: outer nuclear layer. PIS: photoreceptor inner segment. POS: photoreceptor outer segment. RPE: retinal pigment epithelium. BM: Bruch’s membrane. CHO: choroid. Hypopigmented spots and sub-RPE abnormalites were averaged from 4 fundus and OCT images, respectively (n = 4). *P* values were denoted: n.s. *P* > 0.05; \**P* < 0.05; \*\**P* < 0.01; \*\*\**P* < 0.001.

### Increased Sub-RPE Deposits and Thickened Bruch’s Membrane in the Adult DKO Mice

Periodic acid-Schiff (PAS) staining of retinal sections from the adult DKO mice, the CXCR5 KO mice, and C57BL6 WT controls showed the increased BM and the aberrant deposits within the RPE and/or sub-RPE deposits in both 2- and 4-month-old DKO animals, but not in the KO mice and WT controls (Fig. 2A-D). The morphologies of the deposits (punctate and hemisphere) were consistent in shape and location with the hypopigmented spots and sub-RPE abnormalities visualized by fundus examination and OCT images. Quantifications revealed that the numbers of RPE and sub-RPE deposits were significantly higher in the DKO mice (both 2 and 4 months old) than in the KO mice and WT controls (Fig. 2E). BM thickness was also significantly increased in the DKO mice (4 months old) compared to the KO mice and the WT controls (Fig. 2F).

**Figure 2.**
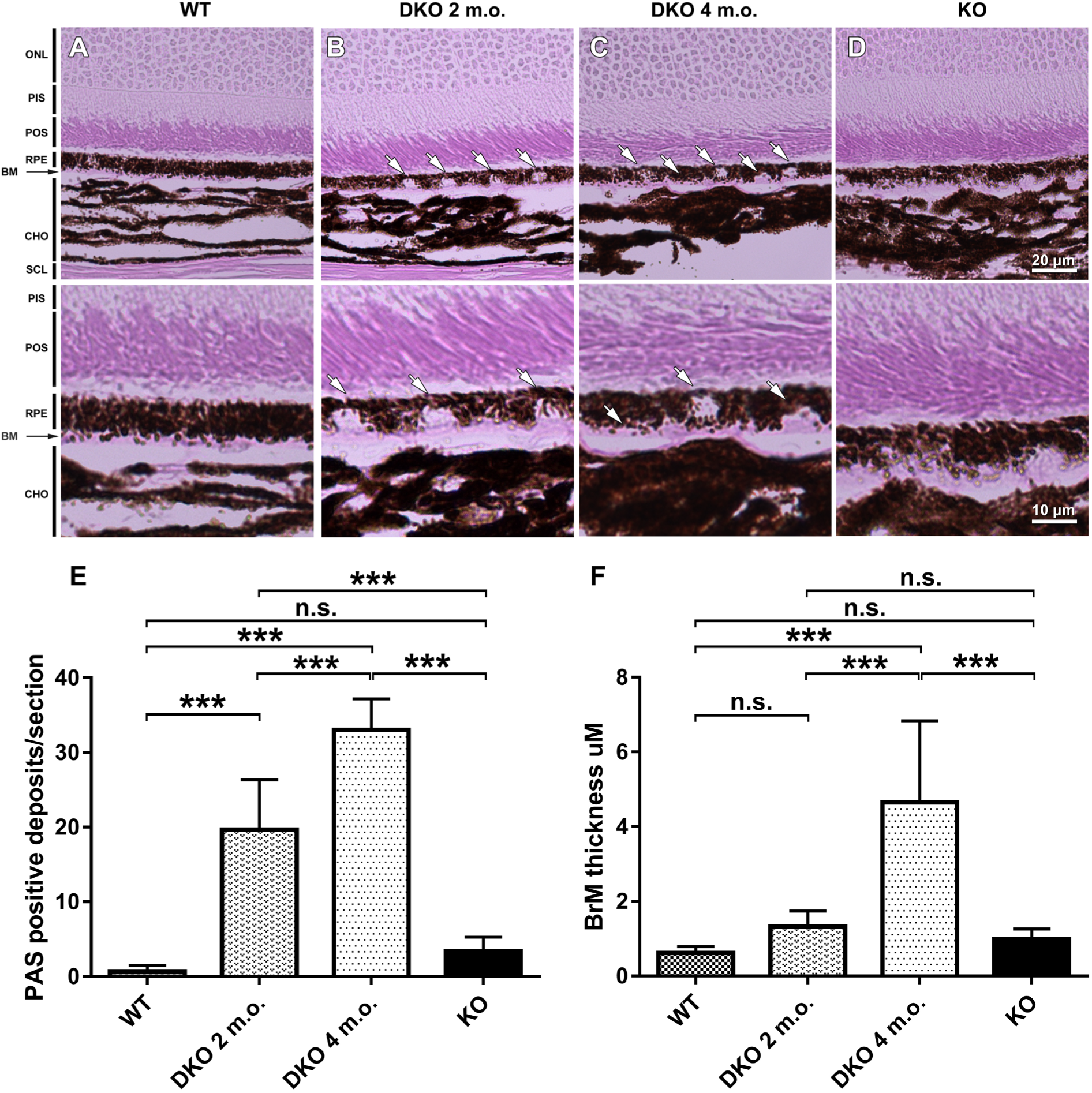
Increased RPE/sub-RPE deposits and thickened Bruch’s membrane (BM) in the adult CXCR5^-/-^.NRF2^-/-^ mice, as demonstrated by PAS staining. Four-month-old C57BL/6 WT mice (A), 2- and 4-month-old CXCR5/NRF2 DKO (B-C), and 4-month-old CXCR5 KO (D) mice histological sections were studied, revealing presence of PAS (+) deposits in RPE-layer of DKO mice at 2-months of age (B); PAS (+) deposits are increasing in size pushing against the thickening Bruchs membrane (visible as pink layer) into choroid layer in 4-months old DKO mice (C). Retinal layers were denoted as follows: ONL, outer nuclear layer; PIS, photoreceptor inner segment; POS, photoreceptor outer segment. RPE, retinal pigment epithelium; BM, Bruch’s membrane; and CHO, choroid. (E) Quantification of PAS (+) deposits within RPE and sub-RPE area. The numbers of RPE/sub-RPE deposits were counted and averaged per group (n = 4). Quantification of the BM thickness (F). The values were averaged from 4 samples per group (n = 4). *P* values were denoted: n.s. *P* p > 0.05; \*\*\**P* < 0.001.

### Increased Autofluorescence and IgG in the Adult DKO Mice

The retinal sections from C57BL6 WT (2 months), CXCR5/NRF2 DKO (2 and 4 months), and CXCR5 KO (4 months) mice were examined for autofluorescence with the 488nm wavelength and endogenous IgG deposits at the interface of the photoreceptor outer segment (POS), RPE, BM, and choroid. Strong spontaneous fluorescence signals were detected on the POS, RPE, BM, and sub-RPE of the DKO mice. Autofluorescence signals were detected in the WT mice and the KO mice as well as DKO mice, but the signal intensity levels were much less in WT and KO mice than in DKO mice (Figs. 3A-3D), indicative of an increased lipofuscin A2E depositions in the DKO mice. IgG levels were increased at the RPE-BM-choroid interface and at the ganglion cell layers of the DKO mice as compared to those of the KO mice and the WT controls (Figs. 3E-H). Western blotting results (without the addition of primary antibody) further confirmed that the endogenous IgG levels were increased in both RPE/BM/choroid and retinal protein extracts (Fig. 3I).

**Figure 3.**
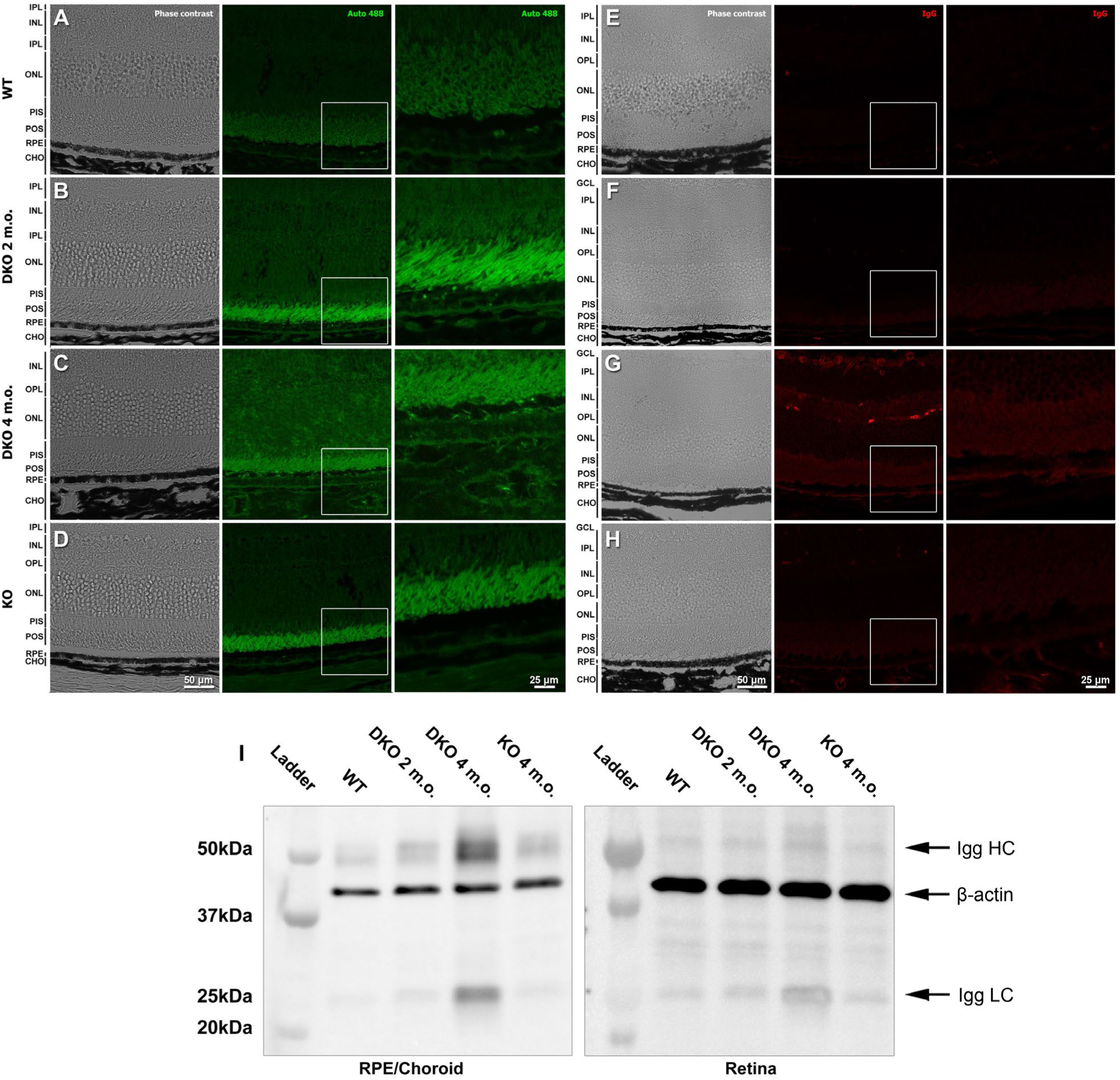
Increased autofluorescence and IgG depositions at RPE and sub-RPE space of the adult CXCR5^-/-^.NRF2^-/-^ mice. Autofluorescence at 488 nm wavelength was examined in the retinal sections prepared from C57B6/J wild type (A, WT, 4 months), CXCR5^-/-^.NRF2^-/-^ (B, DKO, 2 months), CXCR5^-/-^.NRF2^-/-^ (C, DKO, 4 months) and CXCR5^-/-^ (D, KO, 4 months). Endogenous IgG was detected by omitting primary antibody and visualized by anti-mouse IgG secondary antibody conjugated with Cy5 (E-H). Autofluorescence (Alexa 488) and IgG (Cy5) were on the middle column. Phase-contrast images of section morphology shown on the left column. White squares denote the enlarged areas of the corresponding fluorescent images. Retinal layers denoted as follows: GCL: ganglion cell layer. IPL: inner plexiform layer. INL: inner nuclear layer. OPL: outer plexiform layer. ONL: outer nuclear layer. PIS: photoreceptor inner segment. POS: photoreceptor outer segment. RPE: retinal pigment epithelium. BM: Bruch’s membrane. CHO: choroid. (I) The Western blots results of endogenous IgG from RPE/choroid (left panel) and retina (right panel) heavy (HC) and light chains (LC). β-actin used as loading control.

### Increased AMD-Associated Protein Depositions in the Adult DKO Mice

Immunofluorescence (IF) staining to examine the RPE/sub-RPE deposition of the four AMD/drusen-associated proteins—β-amyloid (Aβ), αB-crystallin, apolipoprotein-E (Apo-E), and complement (C5b-9)—revealed that immune reactivity of Aβ and αB-crystallin at the RPE and the sub-RPE space was increased in the 4- and 6-month-old DKO mice but was not detected in the KO mice and the WT controls and was only barely detected in the 2-month-old DKO mice (Fig. 4). In a similar manner, the RPE/sub-RPE depositions of Apo-E and C5b-9 were markedly elevated in the DKO mice compared to the KO mice and the WT control (Fig. 5).

**Figure 4.**
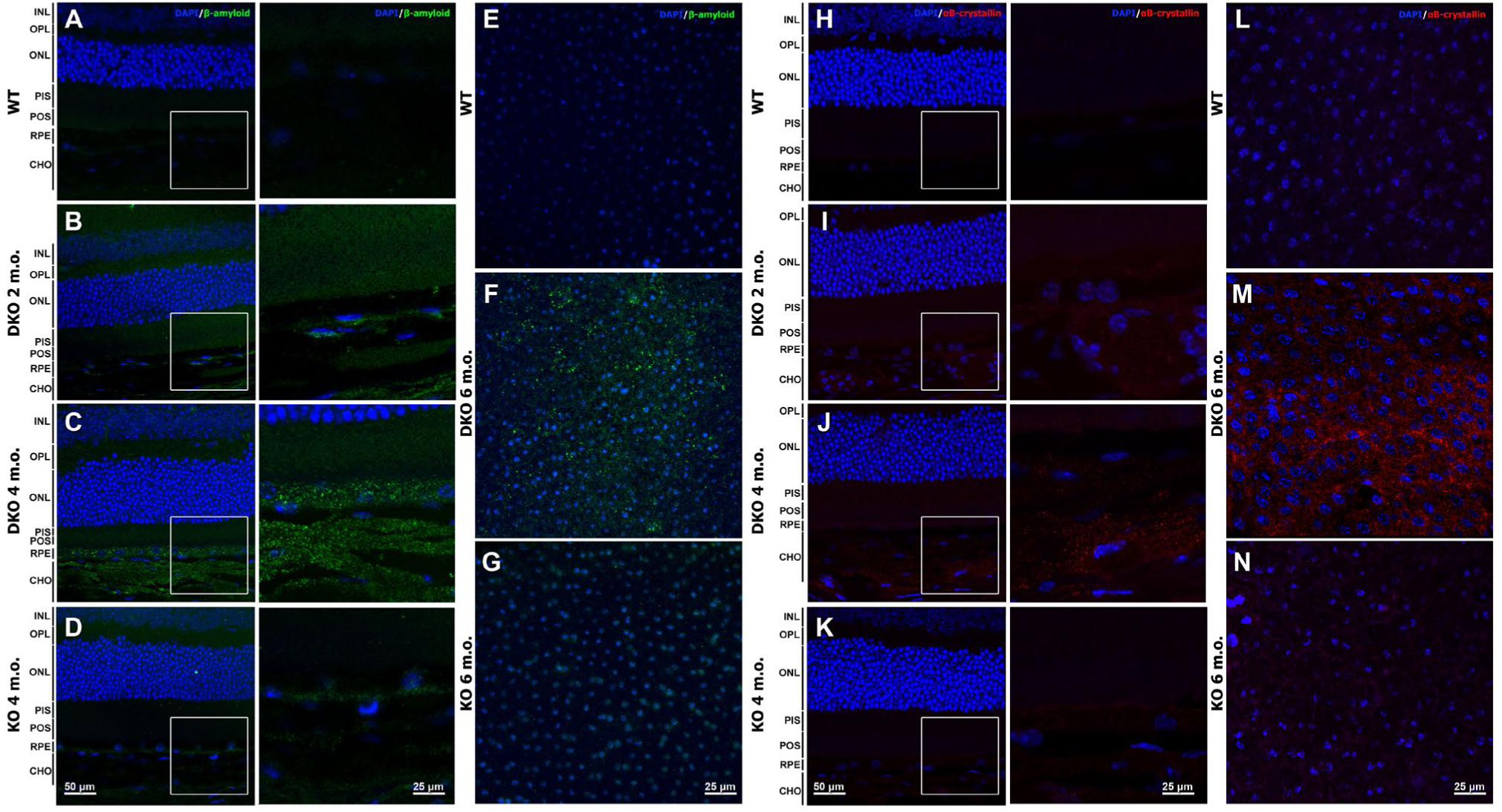
Increased β amyloid and αB-crystallin depositions at the RPE and sub-RPE spaces of the adult CXCR5^-/-^.NRF2^-/-^ mice. Immunofluorescence (IF) staining for β amyloid was performed on the sections prepared from C57B6/J WT mice (A, 2 months); CXCR5^-/-^.NRF2^-/-^ DKO mice(B, 2 and C, 4 months); and CXCR5^-/-^ KO (D 4 months) old mice; as well as on the RPE/BM/choroid/sclera flat mounts of the 6-month-old WT (E), DKO (F) and KO mice. Same groups of sections and flat mounts stained for αB-crystallin (H-N). White squares denote the enlarged areas of the corresponding fluorescent images. Retinal layers were denoted as follows: INL, inner nuclear layer; OPL, outer plexiform layer; ONL, outer nuclear layer; PIS, photoreceptor inner segment; POS, photoreceptor outer segment; RPE, retinal pigment epithelium; CHO, choroid.

**Figure 5.**
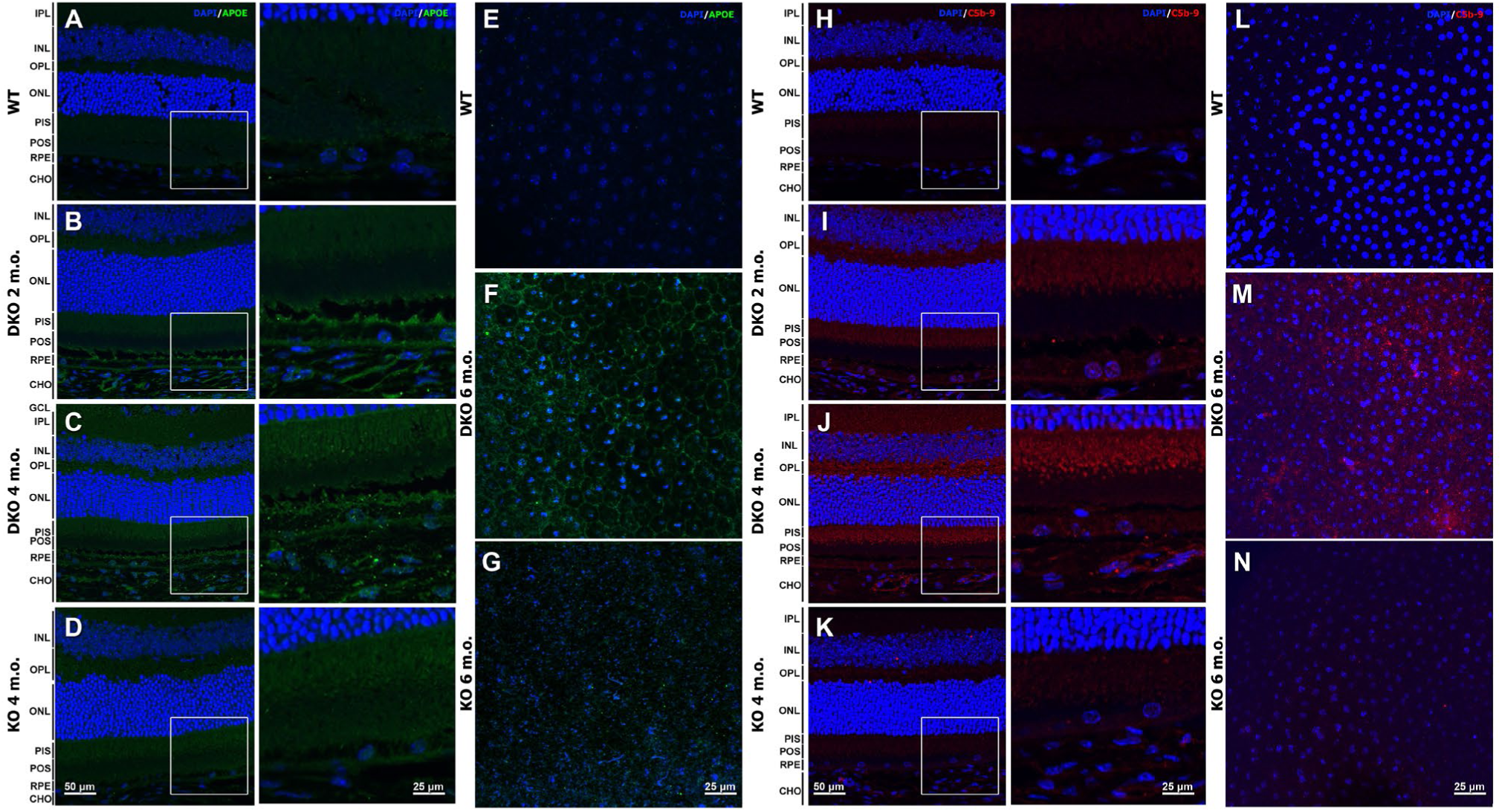
Increased apolipoprotein E and complement 5b-9 depositions at the RPE and sub-RPE spaces of the adult CXCR5^-/-^.NRF2^-/-^ mice. Immunofluorescence (IF) staining for apolipoprotein E (APOE) was performed on the sections prepared from C57B6/J WT mice (A, 2 months); CXCR5^-/-^.NRF2^-/-^ DKO mice(B, 2 and C, 4 months); and CXCR5^-/-^ KO (D 4 months) old mice; as well as on the RPE/Choroid/Sclera flat mounts of the 6-month-old WT (E), DKO (F) and KO mice. Same groups of sections and flat mounts stained for complement 5b-9 (C5b-9) (H-N). White squares denote the enlarged areas of the corresponding fluorescent images. Retinal layers were denoted as follows: GCL, ganglion cell layer; IPL, inner plexiform layer; INL, inner nuclear layer; OPL, outer plexiform layer; ONL, outer nuclear layer; PIS, photoreceptor inner segment; POS, photoreceptor outer segment; RPE, retinal pigment epithelium; CHO, choroid.

### Elevated AMD-Associated Proteins and Reduced RPE Zonula Occludens-1 Protein Levels in Adult DKO Mice

As RPE and sub-RPE depositions of AMD-association proteins could cause RPE degeneration, we, therefore, examined whether the zonula occludens-1 (ZO-1) protein was degraded in the RPE of DKO mice. As indicated by the decreased immune staining signal intensity, ZO-1 was markedly reduced on the RPE flat mounts of the DKO mice compared to the KO mice and the WT controls (Fig. 6). It was also noted that the normal hexagonal shapes of RPE were compromised in the DKO mice (Fig. B), but both the KO mice and the WT controls had a regular layout of ZO-1(+) RPE cells. Western blot results further confirmed that ZO-1 protein levels were reduced in the adult DKO mice (4 and 6 months old) compared to the KO mice and the WT controls (Fig. 6D). Western blot results further revealed that the protein levels of Aβ, Apo-E, C5b-9, and IgG heavy chain (HC) and light chain (LC) were upregulated in the 4-month-old DKO mice and further increased in the 6-month-old DKO mice as compared with the levels in KO mice and the WT controls. Interestingly, the microglia maker TMEM119 was also upregulated in the adults of DKO mice, as compared to the KO and the WT controls.

**Figure 6.**
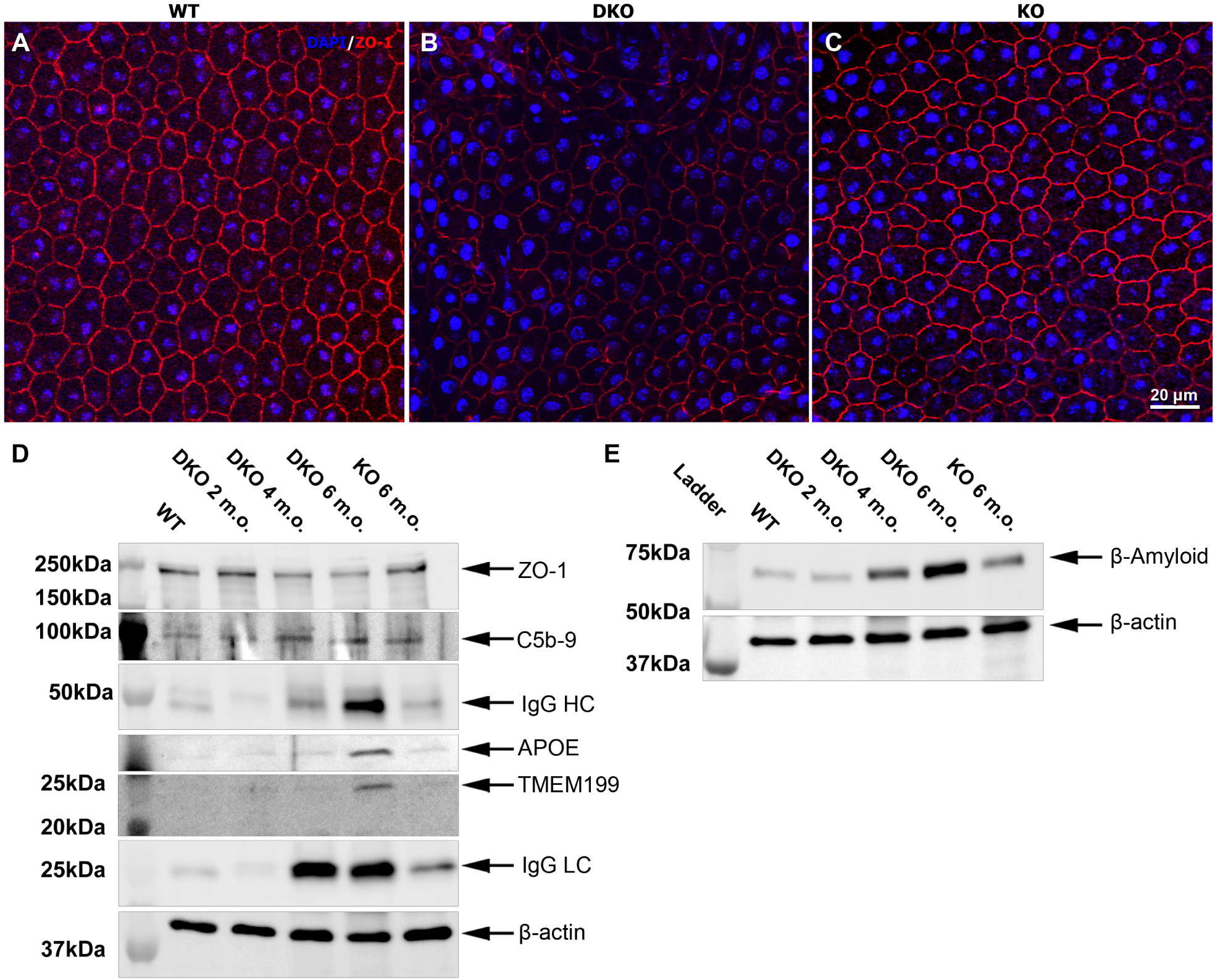
Elevated AMD-associated proteins and reduced RPE zonula occludens-1 (ZO-1) protein levels in the adult CXCR5^-/-^.NRF2^-/-^ mice. Immunostaining for ZO-1 were made in RPE//Choroid/Sclera flat mounts from specimens obtained from 6-month-old C57BL6 WT mice (A), CXCR5^-/-^.NRF2^-/-^ DKO mice (B), and CXCR5^-/-^ KO mice (C). Note that ZO-1 immunostaining intensity of the DKO mice was lower than that of WT and KO mice and that hexagonal-shaped RPE showed abnormalities in the DKO mice (top area in panel B). Western blot analysis of ZO-1; complement 5b-9 (C5b-9); apolipoprotein E (APOE), Transmembrane Protein 119 (TMEM119); heavy (HC) and light chains (LC) IgG; (D) and β-amyloid (E) in RPE/BM/Choroid protein extracts prepared from C57B6/J (WT; 4 months), CXCR5^-/-^.NRF2^-/-^ (DKO; 2, 4, 6 months) and CXCR5^-/-^ (KO; 6 months) old mice. β-actin used as loading control.

### Accelerated Retinal Degeneration and Apoptotic Cell Death in Adult DKO Mice

Finally, we investigated whether retinal degeneration and apoptotic cell death are present in the adult DKO mice. The retinal flat mounts from the 6-month-old DKO mice and the age-matched WT and KO mice were stained with peanut agglutinin lectin, cleaved caspase 3, and microtubule-associated protein 2 (MAP2). Both PNA lectin (+) photoreceptors and MAP2 (+) retinal neurons were significantly decreased in the DKO mice compared with the levels in KO and WT mice (Figs. 7A,7B). Concomitantly, the cleaved caspase 3 (+) cells on the ganglion cell layer and photoreceptor cell layer were markedly increased in the DKO mice compared to the apoptotic cell numbers in retinal specimens from KO mice and WT controls (Figs. 7C, 7D). Further quantification revealed that the photoreceptors and ganglion cell densities were significantly lower in the DKO mice than in the KO mice and the WT controls (Figs. 7E,7F), but the numbers of cleaved caspase 3 (+) apoptotic cells were significantly higher in the DKO mice than in the KO and WT mice. Interestingly slight, but significant increase of caspase 3 (+) was also observed in ganglion and photoreceptor cell layer of adult KO mice (Figs. 7G, 7H).

**Figure 7.**
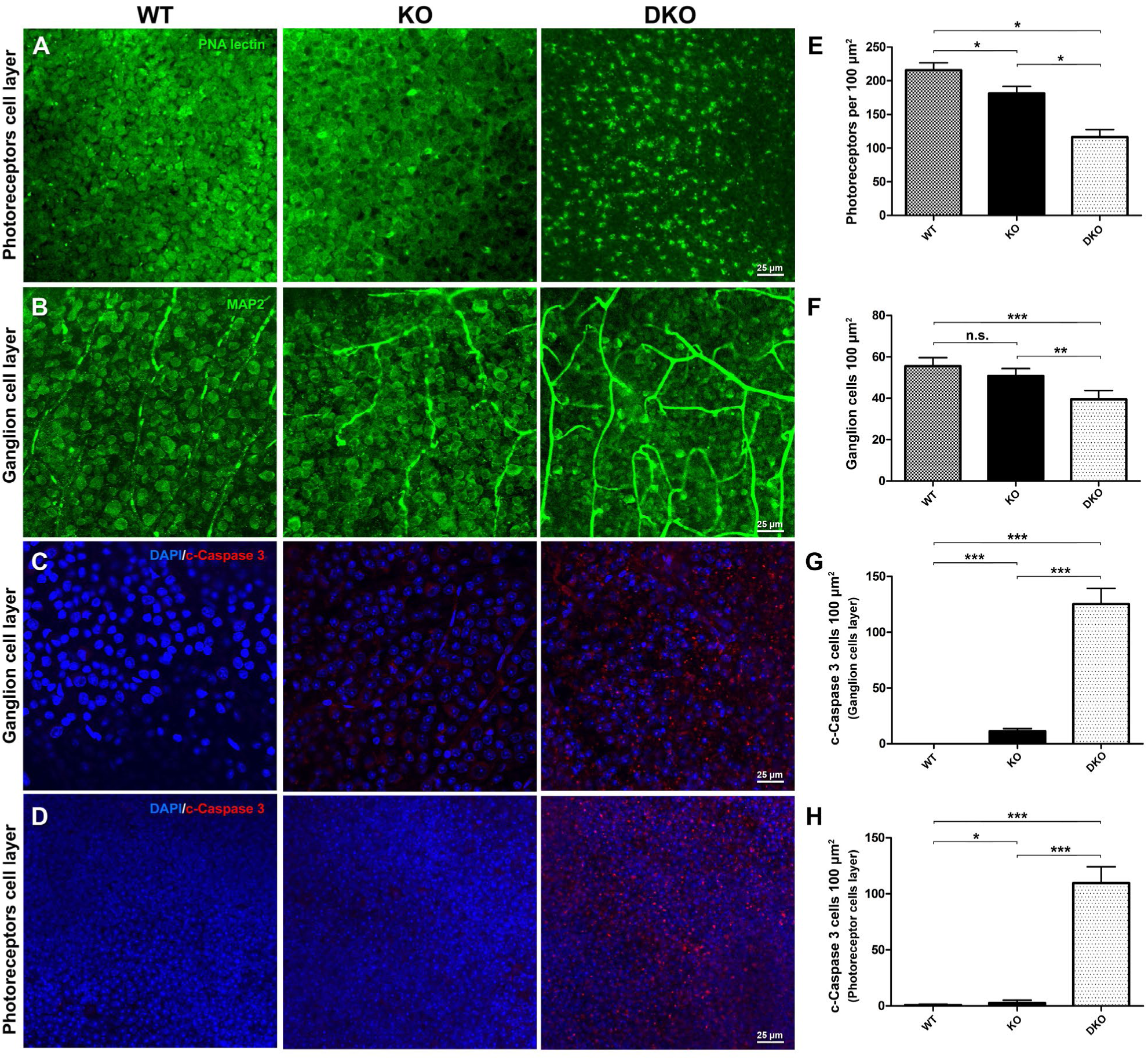
Accelerated retinal degeneration and cell apoptosis in the adult CXCR5^-/-^. NRF2^-/-^ mice. The 6-month-old C5BL6 WT mice, CXCR5^-/-^ KO mice, and CXCR5^-/-^.NRF2^-/-^ DKO mice retinal flatmounts were used for staining and quantification. (A-D) Retinal flatmounts stained with peanut agglutinin (PNA) lectin (A), MAP2 (B), cleaved caspase 3 at the ganglion cell layer (C) and the photoreceptor cell layer depth (D). The quantitative results of PNA lectin (+) photoreceptors (E), MAP2 (+) retinal ganglion cells (F). Caspase 3 (+) ganglion cells layer (G), and Caspase 3 (+) photoreceptor layer depths (H). The values of cell numbers per 100 µm^2^ were averaged from 4 retinal samples (n = 4). *P* values were denoted: n.s. *P*>0.05; \**P* <0.05; \*\**P* <0.01; \*\*\**P* <0.001.

## Discussion

Recently we described retinal degenerative phenotypes in aged Cxc5^-/-^ mice in association with loss of blood-retinal barrier function, the accumulation of AMD-associated proteins such as β-amyloid, Apolipoprotein-E, complement (C3d and C5b-9), and αB-crystallin, and the occurrence of specific autoimmune responses as possible driving forces of retinal degeneration. However, while many of the AMD features are recapitulated in the aged Cxc5^-/-^ mice, after 12 months of life the Cxc5^-/-^ mice start to develop retinal degeneration, which becomes prominent by 18-24 months of age, making it difficult to utilize this animal in studies of potential therapeutic interventions for AMD. Therefore, we tried to create an accelerated animal model of AMD, in which AMD-like phenotypes develop at an early age.

The original conception was that the combination of the two distinct pathological factors (oxidative stress and inflammation/immune deregulation) could accelerate the onset and progression of the AMD pathologies. Initially, we attempted to combine Cxcr5^-/-^ mice with Sod1^mut/mut^ mutant mice (B6SJL-Tg (SOD1*G93A) 1Gur/J; Jackson Laboratory, Bar Harbor, ME, USA), which contain mutant human SOD1 gene (harboring a single amino acid substitution of glycine to alanine at codon 93) driven by its endogenous human SOD1 promoter. While fundus and histological observations revealed that Cxcr5^-/-^.Sod1^mut/mut^ mutant mice did develop some accelerated AMD-like features as early as one month of age (data not shown), their short life span, poor health, low fertility, and small litter size made it challenging to maintain this line, thus reducing the ability to perform complete phenotypic analysis and the potential usefulness of this mouse strain as an AMD model. In addition, as SOD1 mutations are a known factor in amyotrophic lateral sclerosis but are not directly implicated in human AMD, we changed our attention to Nrf2 (the robustly established oxidative stress gene). Nrf2^-/-^ mice are known to be fertile and reasonably healthy.(Zhao, Chen et al. 2011) It is well documented in the literature that Nrf2 is a protective transcription factor controlling the gene expression of a wide range of antioxidants.(Bellezza 2018) Nrf2^-/-^ mice demonstrate typical AMD-like characteristics at 12 months and older due to increased oxidative damage.(Zhao, Chen et al. 2011) Furthermore, the role of Nrf2 has recently been implicated in human AMD and prospected as the therapeutic target.(Bellezza 2018)

The Cxcr5^-/-^.Nrf2^-/-^ mice are viable, healthy, and fertile. With AMD-like features such as BM thickness, aberrant RPE/sub-RPE depositions, increased auto-fluorescence and IgG, elevated complement activation, and retinal cell apoptosis and degeneration occurring at early ages (versus the older ages of single KO of each founding mouse lines), this DKO mouse line could be thought of as an accelerated model for AMD and could be beneficial for studying disease etiology and assessing potential pharmacotherapeutic agents. The AMD-like features detected in the aged Cxcr5 KO, aged Nrf2 KO, and adult DKO mice are summarized in **Table 1**. It is worth noting that we carefully bred out the RD8 mutation from both founder breeding lines and examined the genotyping for the Crb1 gene by Sanger sequencing and subsequently through custom RT-PCR probe, with the assistance of TransnetYX company. While animals on C57BL/6j background usually do not have *rd8* mutation of the *Crb1* gene, it is a common occurrence in animals on C57BL/6N background such as Nrf2^-/-^ mice. The *crb1-rd8* mutation is an emerging problem of retinal degeneration research in mice and was recently addressed by Mattapallil et al. (Mattapallil, Wawrousek et al. 2012)

**Table 1:**
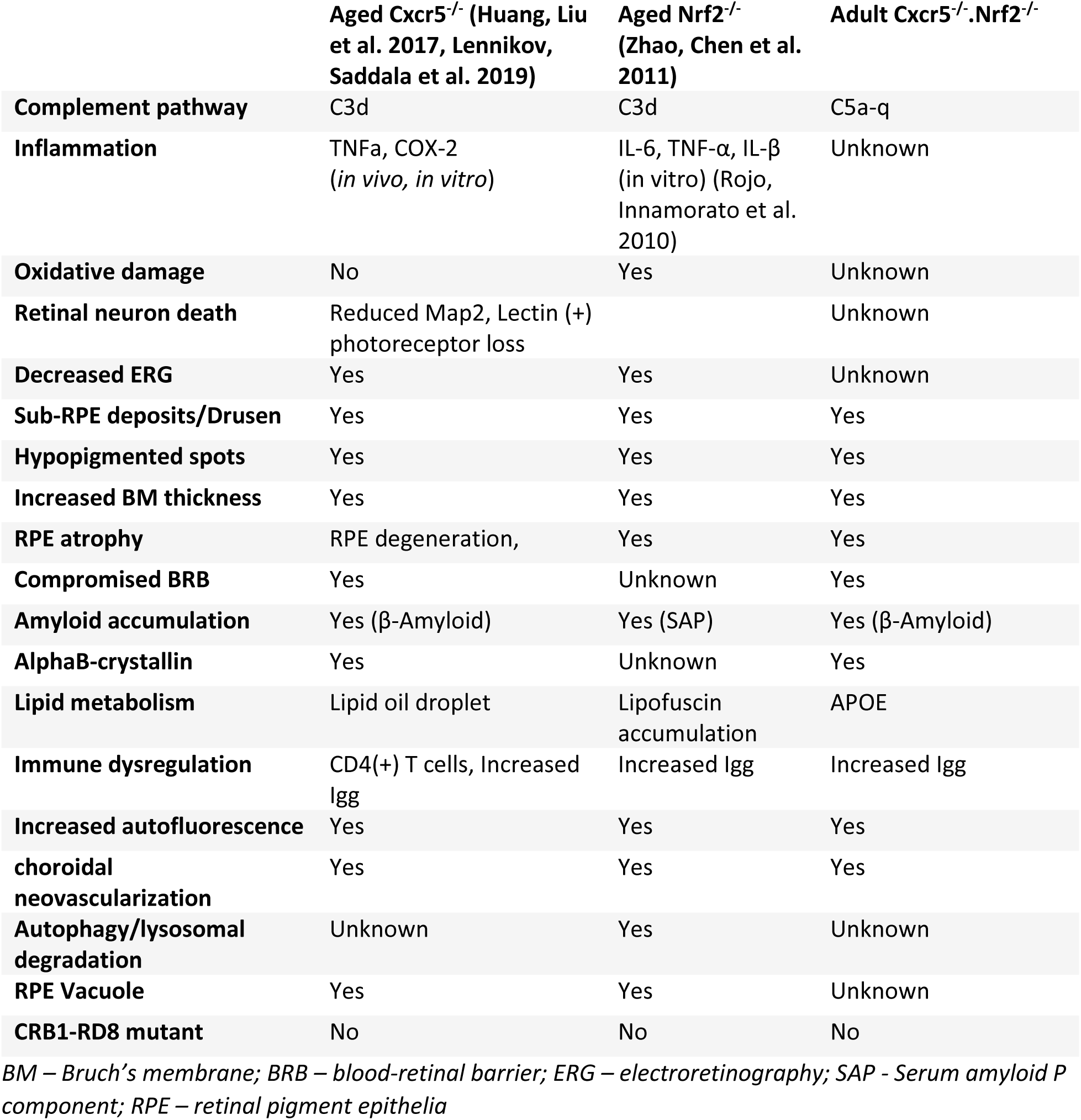
AMD phenotypic feature in aged Cxcr5 KO, aged Nrf2 KO, and DKO.

|  | Aged Cxcr5<sup>-/-</sup> (Huang, Liu et al. 2017, Lennikov, Saddala et al. 2019) | Aged Nrf2<sup>-/-</sup> (Zhao, Chen et al. 2011) | Adult Cxcr5<sup>-/-</sup>.Nrf2<sup>-/-</sup> |
| --- | --- | --- | --- |
| **Complement pathway** | C3d | C3d | C5a-q |
| **Inflammation** | TNFα, COX-2<br>(*in vivo*, *in vitro*) | IL-6, TNF-α, IL-β<br>(*in vitro*) (Rojo, Innamorato et al. 2010) | Unknown |
| **Oxidative damage** | No | Yes | Unknown |
| **Retinal neuron death** | Reduced Map2, Lectin (+)<br>photoreceptor loss |  | Unknown |
| **Decreased ERG** | Yes | Yes | Unknown |
| **Sub-RPE deposits/Drusen** | Yes | Yes | Yes |
| **Hypopigmented spots** | Yes | Yes | Yes |
| **Increased BM thickness** | Yes | Yes | Yes |
| **RPE atrophy** | RPE degeneration, | Yes | Yes |
| **Compromised BRB** | Yes | Unknown | Yes |
| **Amyloid accumulation** | Yes (β-Amyloid) | Yes (SAP) | Yes (β-Amyloid) |
| **AlphaB-crystallin** | Yes | Unknown | Yes |
| **Lipid metabolism** | Lipid oil droplet | Lipofuscin<br>accumulation | APOE |
| **Immune dysregulation** | CD4(+) T cells, Increased<br>Igg | Increased Igg | Increased Igg |
| **Increased autofluorescence** | Yes | Yes | Yes |
| **choroidal**<br>**neovascularization** | Yes | Yes | Yes |
| **Autophagy/lysosomal**<br>**degradation** | Unknown | Yes | Unknown |
| **RPE Vacuole** | Yes | Yes | Unknown |
| **CRB1-RD8 mutant** | No | No | No |

*BM – Bruch's membrane; BRB – blood-retinal barrier; ERG – electroretinography; SAP - Serum amyloid P* *component; RPE – retinal pigment epithelia*

It is certain that increased oxidative stress is a contributing factor to the accelerated process of AMD pathology in the Cxcr5/Nrf2 DKO mice. However, the mechanism through which Cxcr5 deficiency contributes to the development of sub-RPE deposits and AMD pathologies remains to be clarified. Several potential mechanisms may be invovled. Immune deregulation caused by Cxcr5 deficiency may lead to specific autoimmune inflammation in the RPE and the retina, causing RPE defects and AMD development. Our previous studies demonstrated substantial sub-RPE accumulation of IgG, (Huang, Liu et al. 2017) which was later identified as autoantibodies to the AMD-associated proteins, such as annexin-A2, αB-crystallin, ubiquitin-B, and aquaporin-5.(Lennikov, Saddala et al. 2019) Cxcr5 may also be important for the stabilization of microglial cells and the regulation of their response to microenvironment changes. When Cxcr5 was blocked by antibodies in BV-2 cells, microglia showed more activation in response to proinflammatory stimulation.(Lennikov, Saddala et al. 2019) Furthermore, Cxcr5 may play a role in the maintenance of RPE homeostasis and polarization states, particularly in stress conditions such as aging, and oxidative and pro-inflammatory insults. Whether and how these mechanisms interplay to exacerbate the AMD pathological process in Cxcr5/Nrf2 DKO mice remains to be determined.

Among the perturbed biological pathways and processes in AMD are complement pathways, cytokines, chemokines, phagocytosis, autophagy, lipid metabolism, and oxidative stress, many of which have been investigated to generate animal models of AMD that mimic the AMD features observed in humans. Although mice are popularly used as an AMD model, AMD-like pathologies often develop at the advanced age,(Rakoczy, Zhang et al. 2002, Ambati, Anand et al. 2003) and in some cases require additional stimuli, such as feeding with the high-fat diet, exposure to cigarette smoke, blue light,(Cousins, Espinosa-Heidmann et al. 2002) or injection by lipid oxidants.(Hollyfield, Bonilha et al. 2008, Baba, Bhutto et al. 2010) To overcome this issue, we sought to generate an accelerated mouse model of AMD by crossing Cxcr5 KO mice and Nrf2 KO mice and found that these DKO mice develop the AMD phenotypic features at earlier ages (2 to 6 months) than those aged models (9 months and older). These new DKO mice may provide to be suitable for the elucidation of the underlying mechanisms of AMD pathology and the evaluation of new treatment strategies and drug candidates for AMD.

## Materials and Methods

### Animals, Genotyping and Breeding Strategy

All experiments were approved by the University of Missouri Institutional Animal Care and Use Committee (protocol number: 9520) and performed in accordance with the “Statement for the Use of Animals in Ophthalmic and Vision Research” of the Association for Research in Vision and Ophthalmology. The [B6.129S2(Cg)-Cxcr5tm1^Lipp/J^] (CXCR5, KO), B6.129X1-Nfe2l2^tm1Ywk^/J (NRF2) and [C57BL/6J] (WT) mice strains were purchased from Jackson Laboratory. CXCR5 mice (https://www.jax.org/strain/006659) and NRF2 mice (https://www.jax.org/strain/017009) are on a mixed C57/BL6J/N background with the predominance of C57/BL6J genome. All mice were housed at the special pathogen-free animal facilities of the Bone Life Sciences Center at the University of Missouri and were fed normal chow diets and provided with water *ad libitum*.

The two founder mice were bred to generate the Cxcr5/Nrf2 double knockout (DKO) mice. The RD8 mutation in the Crb1 gene incorporated in the Nrf2^-/-^ founder animal was bred out to achieve the stable Cxcr5/Nrf2 (DKO) mouse line with wild-type RD8 genotype (Cxcr5^-/-^.Nrf2^-/-^.Crb1-Rd8^wt/wt^). The description of the breeding program is presented in Suppl. Fig.1. The Cxcr5/Nrf2 DKO mice were maintained and inbreed-crossed to give birth to more progenies for further research.

Genotyping was performed with the assistance of Transnetyx (Cordova, TN, USA), Outsourced PCR Genotyping Services (www.transnetyx.com) by real-time polymerase chain reaction (PCR) genotypic assay. Genomic DNA (50 ng) was extracted from tail tips and amplified using custom-designed genotyping primers (TransnetYX). Animals were validated for knockout of the CXCR5, NRF2 gene, the presence of the neomycin resistance, and LacZ genes. All mice were screened for the presence of Rd8-associated nucleotide deletion, using the Rd8 genotyping probe designed by Transnetyx, based on our previous Sanger sequencing data of the region 3600-3700 of the Crb1 gene (canonical transcript M_133239). (Lennikov, Saddala et al. 2019)

### Anesthesia and Euthanasia

During *in vivo* experiments, mice were anesthetized by intraperitoneal injection of ketamine hydrochloride (100 mg/kg body weight) and xylazine (4 mg/kg body weight) at the experimental endpoints of 2, 4 and 6 months of age. For tissue collection, mice were euthanized by intraperitoneal injection of ketamine hydrochloride (300 mg/kg body weight).

### Fundus Examination and Optical Coherency Tomography

Mice were anesthetized. Pupils were dilated with 1% tropicamide (Sandoz, US). The cornea was protected with (hypromellose ophthalmic demulcent solution) Gonak 2.5% (Akorn LLC, Akorn, OH, USA) transparent gonioscopy gel. Fundus examination and optical coherence tomography (OCT) was performed with a Micron IV retinal imaging microscope system (Phoenix Research Labs, Inc., Pleasanton, CA, USA). Grayscale OCT images were further processed to produce the heatmap images, where grayscale range 0 (black) to 255 (white) colors were assigned using Photoshop CC gradient map function (Adobe, San Jose, CA, USA). Fundus hypopigmented spots and OCT abnormalities were quantified by “masked” observer.

### Confocal Microscope Imaging

Visible light images were acquired using the EVOS FL Color microscope (Thermo Fisher Scientific, Waltham, MA, USA). Fluorescent images were acquired with a LeicaSP8 laser confocal microscope (Leica AG, Wetzlar, Germany).

### Histology and Immunofluorescent Analysis

The eyeballs were fixed with HistoChoice Molecular Biology fixative (H120-4L, VWR Life Science, Radnor, PA, USA) for 12 hours and stored in phosphate-buffered saline (PBS; 10010023, Thermo Fisher Scientific) until specimens were processed for paraffin embedding and sectioning (5 µm thick) and stained by periodic acid and Schiff reagents staining, as reported previously.(Lennikov, Saddala et al. 2019) Sections intended for autofluorescence evaluation were deparaffinized, rehydrated and mounted without staining. Sections for the detection of autoantibodies accumulation were blocked and permeabilized with 0.5% Triton X-100 (85111, Thermo Fisher Scientific) and blocked with 2.5% bovine serum albumin solution (BSA; A7906-50G, Sigma-Aldrich, St. Louis, MO, USA) for 1 hour at room temperature and incubated overnight at 4 °C with antimouse (ab6563) secondary antibody.

The primary antibody used for immunofluorescent analysis in rehydrated sections included: beta-amyloid (36-6900; 1:100, Thermo Fisher Scientific); αB-crystallin (Ab151722, 1:50; Abcam, Cambridge, England); ApoE (AB947, 1:50; MilliporeSigma, Burlington, MA, USA); Complement 5b-9 (204903-1MG, 1:50; EMD Millipore Corp). Following PBS-Tween 20 0.05% (PBS-T) washing. The immune reactive signals were visualized by Cy5 conjugated anti-rabbit (ab97077, Abcam), anti-mouse (ab6563, Abcam) and anti-goat (ab150131, Abcam) secondary antibody 1:1,000 (Abcam). Sections were counterstained with 4′, 6-diamidino-2-phenylindole (DAPI) 1:5,000 (Sigma-Aldrich) and mounted with ProLong Diamond antifade reagent (P36961, Thermo Fisher Scientific).

### Retinal and RPE–Choroid-Sclera Complex Flat Mounts

Mouse retinas were fixed and isolated as reported previously. (Lennikov, Saddala et al. 2019) Briefly, under a dissection microscope, the anterior segment tissues, vitreous, and were removed to produce an eyecup. Then retina was gently separated from RPE–choroid sclera complex (RCSC), and four relaxing radial incisions were made to RCSC and retina in order to produce the flat-mount. The retinas and RCSC was blocked and permeabilized with a solution composed of 2.5% bovine serum albumin (BSA; 00000) in PBS overnight with 0.01% Triton-X; RCSC samples were then incubated with beta-amyloid (36-6900; 1:100; Thermo Fisher Scientific), αB-crystallin (Ab151722, 1:50; Abcam), ApoE (AB947, 1:50; MilliporeSigma); Complement 5b-9 (204903-1MG, 1:50; EMD Millipore Corp); and anti-ZO-1 (1:100, 402200, Thermo Fisher Scientific,). Retina samples were incubated for 24 hours at 4° C with gentle agitation, in peanut agglutinin (PNA) lectin (1:50, L32460; Thermo Fisher Scientific), MAP2 (1:100, 13-1500; Thermo Fisher Scientific), and cleaved-caspase 3 (1:50, D175, 5A1E; Cell Signaling Technology, Danvers, MA, USA); before being washed three times for 10 min with PBS-T; PNA lectin-stained retinas were counterstained with 4′, 6-diamidino-2-phenylindole (DAPI) 1:5,000 (Sigma-Aldrich) and mounted with ProLong Diamond antifade reagent (Thermo Fisher Scientific). Remaining samples were incubated for 24 hours with Cy5-conjugated anti-rabbit (ab97077), anti-mouse (ab6563) and anti-goat (ab150131) secondary antibody (1:1,000; Abcam) and counterstained with DAPI 1:5,000 (Sigma-Aldrich). After another PBS-T washing three times for 10 min, the samples were mounted on slides with ProLong Diamond antifade reagent. Samples incubated with a blocking buffer (primary antibody was omitted), followed by secondary antibody incubation were used as the background control.

### Western Blot Analysis

Retinal and RCSC were isolated on ice, and lysates were prepared as previously described. (Lennikov, Saddala et al. 2019) Thirty micrograms of total proteins were separated by SDS-PAGE gel (Mini-Protean Precast Acrylamide Gels, Bio-Rad, Hercules, CA, USA) and further transferred to the nitrocellulose membrane (Trans-Blot Turbo transfer pack, Bio-Rad). Membranes were blocked with 2.5% BSA and incubated with primary antibodies: beta-amyloid (36-6900; 1:100, Thermo Fisher Scientific); anti-ZO-1 (1:1000, 402200, Thermo Fisher Scientific), anti-C5b-9 (204903, 1: 1000, EMD Millipore Corp); anti-ApoE (AB947, 1:1000, EMD Millipore Corp); Transmembrane Protein 119 (TMEM119) (1:1000, ab209064, Abcam) or β-actin (PA1-21167; 1:2,000; Thermo Fisher Scientific).

The target protein bands were detected with a horseradish peroxidase (HRP)–conjugated antibody (170-6515, 172-1011, 1720-1011, 1:1,000; Bio-Rad), which was visualized by chemiluminescence with Clarity Western ECL substrate (Bio-Rad) and imaged using the LAS-500 Imaging System (General Electric, Boston, MA, USA). The resulting band sizes were resolved using Precision Plus Protein™ Kaleidoscope™ Prestained Protein Standard (1610375, Bio-Rad). For autoantibody detection, the Western blot on RCSC and retinal lysates were separated as described above. Following blocking, membranes were incubated with Goat Anti-Mouse IgG (Heavy + Light)-HRP conjugate (170-6516, 1:1,000; Bio-Rad) overnight and following PBS-T washing detected using the LAS-500 Imaging System (General Electric).

### Statistical Analysis

All experiments were performed in triplicates. Experimental values were expressed as the mean ± standard deviation (SD) for the respective groups. Statistical analyses were performed with GraphPad Prism software (https://www.graphpad.com/scientific-software/prism/). A one-way ANOVA with Tukey multiple comparisons was used. A p-value of less than 0.05 was considered significant. The following designations for the *P*-value were as follows: n.s. *P* >0.05; * *P*<0.05; ** *P* <0.01; *** *P* <0.001.

## Declarations

## Acknowledgments

The authors would like to acknowledge the following contributors: Allen Raye (University of Missouri Department of Biomedical Sciences, Columbia, Missouri, USA) and Lijuan Fan (University of Missouri, Columbia, Missouri, USA) for assistance with animal resources; Molecular Cytology core (University of Missouri, Columbia, Missouri, USA) for technical assistance with confocal imaging; Ms. Sharon Morey (University of Missouri, Department of Opthalmology Columbia, Missouri, USA) for editing the manuscript; Ms. Catherine Brooks J. (University of Missouri, Department of Opthalmology Columbia, Missouri, USA) for “masked” quantification and additional language corrections.

## Availability of data and materials

All data generated and analyzed in the current study are included in this published article and its supplementary information. Breeding pairs of Cxcr5^-/-^.Nrf2^-/-^.Rd8^wt/wt^ mice can be provided upon reasonable request to the corresponding author.

## Authors’ Contributions

The study was conceived and designed by H.H. and A.L.; H.H. and A.L. performed animal breeding and genotyping; H.H. and A.L. performed *in vitro* and *in vivo* experiments and evaluations. The manuscript was written by H.H. and A.L. and critically revised by H.H. Both authors reviewed and accepted the final version of the manuscript.

## Ethics approval

All experiments were approved by the Institutional Animal Care and Use Committee of the University of Missouri School of Medicine (protocol number: 9520) and were in accordance with the guidelines of the Association for Research in Vision and Ophthalmology Statement for the use of animals in ophthalmic and vision research.

